# Creation and validation of LIMÓN - Longitudinal Individual Microbial Omics Networks

**DOI:** 10.1101/2025.03.18.644048

**Authors:** Suzanne Alvernaz, Beatriz Peñalver Bernabé

## Abstract

Microbial communities are dynamic structures that continually adapt to their surrounding environment. Such communities play pivotal roles in countless ecosystems from environmental to human health. Perturbations of these community structures have been implicated in disease processes such as Crohn’s disease and cancer. Disturbances to existing ecosystems often occur over time, making it essential to have robust methods for detecting longitudinal alterations in microbial interactions as they develop. Existing methods for identifying temporal microbial community alterations have focused on abundance alterations in individual taxa, rather than relationships between the taxa, known as microbial interactions. Identifying these interactions overtime provides a fuller understanding of how the microbial ecosystem changes as a whole. To fill this gap, we have developed a pipeline that handles the complicated nature of repeated compositional count data, LIMÓN – Longitudinal Individual Microbial Omics Networks. This novel statistical approach addresses key challenges of modeling temporal and microbial data including overdispersion, zero-inflated count data, compositionality, repeated measure design sample covariates over time, and identification of individualized or sample specific networks. This approach allows users to denoise covariate effects from their data, return networks per time point, identify interaction changes between each time point, and return individual networks and network characteristics per sample/time point. In doing so, LIMÓN provides a platform to identify the relationship between network interactions and sample features of interest over time. Here we show LIMÓN, in simulation studies, can accurately remove covariate effects, render sample specific networks, and better recover underlying network edges from covariate confounded data. Analysis of a longitudinal infant microbiome and diet dataset illustrates LIMÓN’s novel utility to identify key microbial interactions related to diet type across time.

**AUTHOR SUMMARY:** Microbes (bacteria, fungi etc.) are integral components of many ecosystems, from the environment to the human body, where they can shift between healthy and disease states. Microbes do not exist alone but in rich diverse communities. Yet, many current methods used to study microbe alterations in disease focus on changes in individual microbes rather than how the entire community adapts. To better understand how microbial communities shift, we developed an open-source tool called LIMÓN, which allows users study how these relationships shift over time. By leveraging robust statistical techniques, LIMÓN can account for the complexities of the data, such as covariates and differences between individual samples. This approach helps us uncover important patterns in how microbes interact in various conditions. In this example, we applied LIMÓN to data from infants who were fed three different diets during the first year of life and identify specific microbial interactions related to diet that change overtime. This work broadens the scope for exploring microbial ecosystem dynamics in health and nature, offering a more comprehensive perspective vs traditional method used.

## INTRODUCTION

Microbial communities shape the world we live in. Bacteria, archaea, and fungi form complex communities that support all aspects of life from human health (1) (2) (3) (4), marine environments (5), and bioremediation processes (6). Interactions among microbes are very complex and dynamic with many potential biological mechanisms for observed co-occurrence patterns (mutualism, commensalism etc.) (7) Further, microbial communities are unique to the individual subject or sample site (8). This makes them sensitive to external influences or sample-specific conditions. Host characteristics can change over time and impact microbial community structure and hence its functions. For instance, gut microbial abundances and functions are known to oscillate with host circadian rhythms (9) (10). Antibiotic use during a longitudinal cohort study can confound true underlying associations between microbial communities and study outcomes (11). Sex differences and hormonal changes have also been identified as being associated with different microbial community phenotypes (12). Therefore, there is a need for methods that can model dynamic processes while eliminating confounding sample characteristics that may mask the real interactions between the members of the microbial communities and reveal the distinctions at the individual level.

Interactions between the members of microbial communities can be quantified as networks with the nodes as the microbial organisms and the edges representing interactions between community members (13). Many methods for inference of microbial networks exist including those based on correlations, (14), regularized linear regression (15), or conditionally dependent graphs (16), to name a few. Overdispersion, or higher than expected variability in a dataset, is common in microbial data as well as the zero-inflated count/compositional nature of the data which must be addressed when inferring graphical networks (17). Methods such as SparCc (18), FastSpar (19) and SParse InversE Covariance Estimation for Ecological ASsociation Inference (SPIEC-EASI) (16) have been developed to account for the zero-inflated and compositional nature of microbial data (20) and are highly utilized, user friendly pipelines for developing microbial networks in cohort studies with sparse data. Yet, these network methods cannot handle longitudinal data or render individualized networks.

Since microbial communities adapt to changes in environment, methods to assess their interactions longitudinally and individually are essential but still underdeveloped. Some methods are available to investigate temporal interactions in microbial communities such as extended local similarity analysis (21), the theoretical-based generalized Lotka-Volterra (22), regularized S-map (23), and dynamic Bayesian networks (24). Yet, these approaches may require large sample sizes; some fail to adequately account for the compositional nature of microbial data; or are unable to associate a specific (temporal) interaction between community members with a feature of interest (network rewiring) (25) (26). MAGMA is one current method to determine microbial communities while adjusting for sample covariates (27) and Kuijjer and colleagues have recently developed a method for estimation of individual networks based on gene expression data, Linear Interpolation to Obtain Network Estimates for Single Samples (LIONESS) (28). Melograna and colleagues recently combined these tools, MAGMA and LIONESS, to generate individualized microbial networks to study IBD (29). However, their approach has several limitations: first, it does not control for within-subject error, which is necessary in longitudinal data as it involves repeated sample measures (27); second, it cannot include temporal covariates (e.g., weight change in infant studies); and finally, the network inferenced used does not account for the zero-inflated and compositional nature of microbial data when constructing their networks. A model specific to microbial data that allows users to estimate community interactions over time, account for perturbations at an individual level, and relate these dynamics to their study features of interest by creating individualized networks is missing.

To fill this gap, we have developed a novel computational pipeline, LIMÓN – Longitudinal Individual Microbial Omics Networks. LIMÓN is a user-centered R package to analyze longitudinal microbial cohort data at a personalized level. LIMÓN overcomes several unique difficulties of modeling temporal sample-specific microbial community interactions, including overdispersion, zero-inflated count data, compositionality, temporal covariates, repeated measure study design, and returns network characteristics at an individual sample level. We do so by employing mixed effect models with a user-specified distribution (e.g., linear, negative binomial, zero-inflated negative binomial), to extract the covariate denoised counts, and then inferring networks per time point with SPIEC-EASI, which can adequately account for the zero-inflated and compositional nature of microbial data. LIMÓN the provides an estimation of individual networks or networks per sample per time point by including SPEIC-EASI within LIONESS (28) to account for zero-inflated compositional data when inferring interactions between microbial species. Finally, the built-in statistical inference functions allow users to identify key microbial interactions related to their feature of interest overtime. In this work, we describe LIMÓN and validate its accuracy to return single sample estimates for microbial networks. We then apply LIMON to identify longitudinally altering network edges in an infant microbiome dataset of different diet types.

## METHODS

### A. LIMÓN Design

LIMÓN is designed for discrete dynamic measurements (**Fig. 1**). Below we describe LIMÓN procedures in detail.

**Fig 1.**
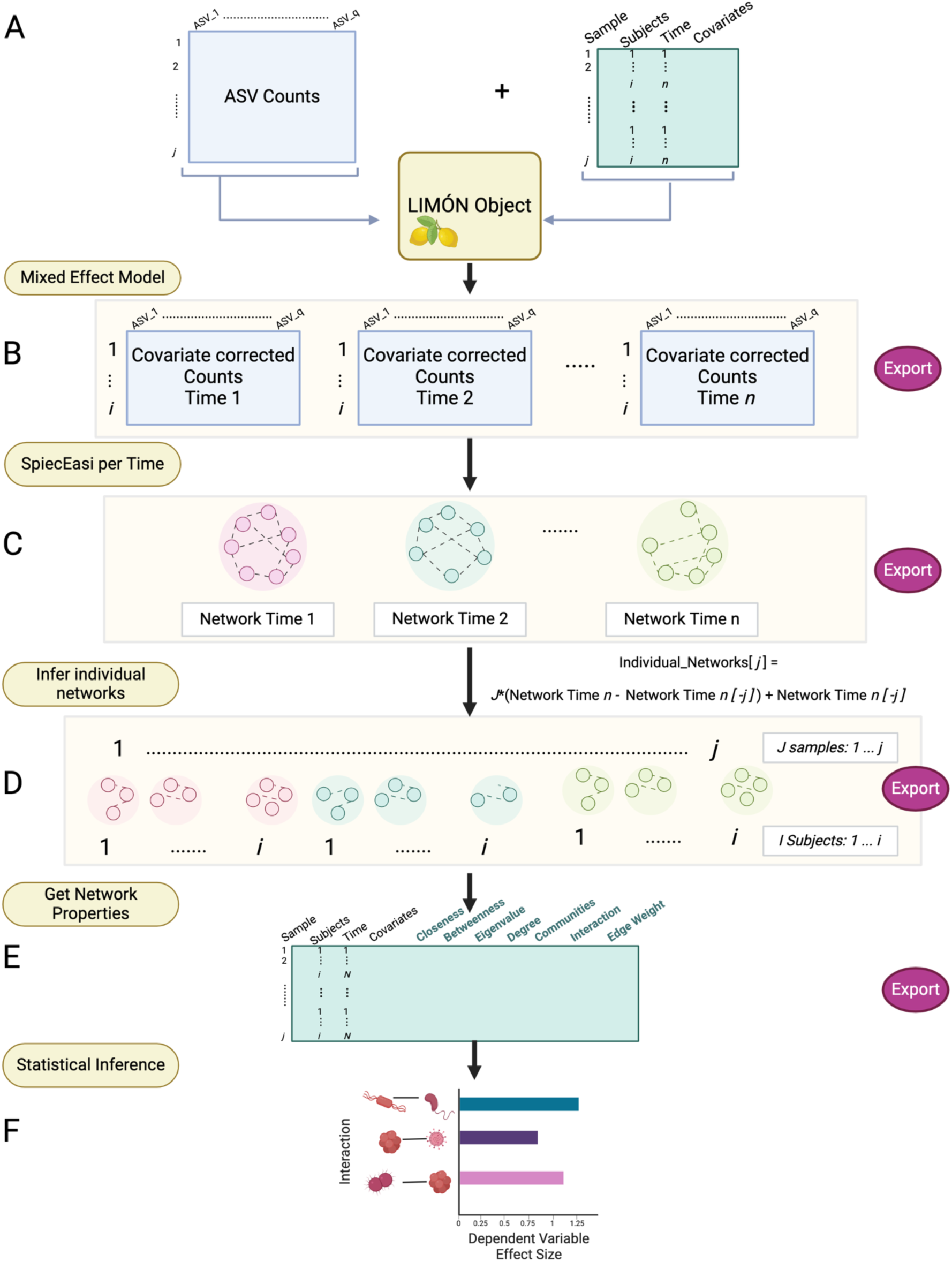
Graphical Representation of LIMÓN workflow and outputs. (A) Denoised microbial counts are combined with sample metadata. (B) Counts are corrected with a mixed effect model to remove variability due to covariates. (C) SPIEC-EASI co-abundance networks per time point are created with denoised counts. (D) Individual networks are inferred, and network characteristics are calculated. (E) Network properties are extracted per sample. (F) statistical inference to identify which interactions are related to the study feature of interest. Created in https://BioRender.com.

#### A1. Data structure

A sample designates a set of microbial counts (taxa or genes) for subject *i* (*i* ∈ [1, *I*]) at time point *n* (*n* ∈ [1, *N*]) for a total of *J* samples (*j* ∈ [1, *J*], *I* subjects and *N* time points). An unnormalized count table, such as amplicon sequencing variants (ASV), of *Q* microbial taxa [(*q* ∈ [1, *Q*]) with *J* total rows (*jth* row = subject *i* at time point *n*) is merged with a user specified sample dataset with *C* covariates [(*covariates* ∈ [1, *C*]) and any features of interest using the *LIMON_Obj* function. (**Fig. 1A**). Users can alternatively use the *Phyloseq_to_LIMON* function to create the initial LIMÓN object from an existing phyloseq object (30).

#### A2. Covariate correction

To correct for sample covariates and optionally zero-inflated over dispersed data, each ASV *q* is fit separately to a mixed effect model with a user specified distribution and subjects (*i* ∈ [1, *I*]) as random effects (varying intercept) using the *LIMON_DistrFit* function. As an example, to remove one subject-level covariate (*F*_1_) with a normal linear mixed effect model option this can be modeled as,

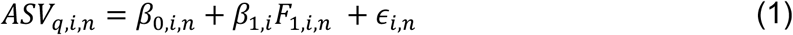

where *ASV_q,i,n_* is the original observed value of ASV *q* for subject *i* at time *n*; *F*_1,*i,n*_ is the covariate 1 value for subject *i* at time point *n*; *β*_0,*i*_ is the estimated intercept; *β*_1,*i*_ is the effect of covariate 1 on ASV *q* for subject *i*, and ∈_*i,n*_ is the error. *β*_0,*i*_ is defined as

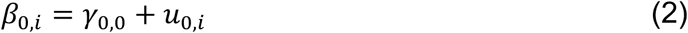

where *γ*_0,0_ is the common fixed intercept value across all individuals and *u*_0,*i*_ is an estimate of the individual intercept deviation for subject *i*, thus allowing variation of the intercept value of ASV *q* per subject, thus controlling for subject specific error. Covariates are modeled at the subject level, shown in equation 3, where *γ*_1,0_ is the fixed effect of *covariate*_1_ across all subjects.

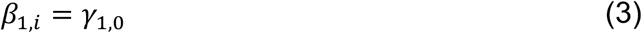

To increase flexibility, users have the option to specify three different distributions: 1) a negative binomial model; 2) zero-inflated Gaussian model; or 3) zero-inflated negative binomial mixed effect model, all based on the *NBZIMM* package in R (31). When deciding which mixed effect model to use, users should determine the distribution that best fits their dataset. For a linear model, the *lmer()* function from the *lme4* R package is employed with default parameters and the intercept varying per subject (32).

From the mixed model output, denoised ASV counts are extracted by subtracting the estimated covariate effect from the original observed values as shown in equation 4.

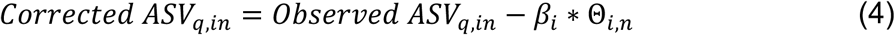

Here, Θ_*i,n*_ is the covariate value for each subject *i* at each time point, *n*. For the one covariate example shown in equation 1, the denoised value of taxa *q* would be estimated as follows (eq. 5).

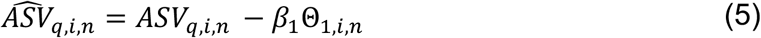

Once denoised, the corrected taxa values are brought above 0 by adding the minimum value and are rounded back to integers. Adjusted counts are then separated into count tables for each time point, *n* (**Fig. 1B**). It is important to note that the adjusted counts have the time variance in the count data preserved, but the noise from the specified (temporal) covariates is removed.

#### A3. Compositionality and network inference per time point

Next, the co-abundance networks from each time point from the denoised ASVs are determined using SPIEC-EASI (**Fig. 1C**, *LIMON_NetInf_Time* function). SPIEC-EASI leverages either a graphical LASSO (33) or a neighborhood selection framework (34) for network inference. Denoised taxonomical counts are centered-log-ratio transformed by SPIEC-EASI (16) (35). The function *LIMON_Edge_Networks* renders the edge table of the computed co-abundance network and *igraph* object (36) per time point *n*. The optimal covariance matrix for each network is extracted using SPIEC-EASI defaults. The diagonal elements are set to 0 and the edges are filtered by a user specified absolute threshold for that optimal covariance matrix (default = 0.2). The printed graphs in R are by default undirected *igraph* plots with edges colored by sign of the interaction. An edge table is stored for each time point.

#### A4. Single Sample (Individual) Network Estimates

Next, LIMÓN infers a network for each *jth* sample (subject *i* at each *n* time point) using a modified version of LIONESS (28) that allows the determination of the co-abundance microbial network using SPIEC-EASI to account for zero-inflated and compositional data (**Fig. 1D**). This method calculates the *jth* sample-specific network by considering the effect of the removal of the sample *j* in the co-abundance network structure in the following manner:

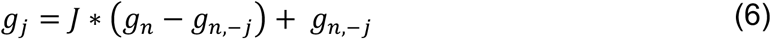

where *g_n,–j_* is the covariance network of all samples except sample *j* at time point *n*. All *J* individualized networks are stored in the LIMÓN output object. From these, the *LIMON_Centralities* function will calculate network centralities of mean betweenness, mean closeness, average node degree, network eigenvalue, and total number of communities (37) per individual network *j* at time *n* and store them in the original sample data frame. Finally, all unique taxa to taxa interactions are extracted as described in **A3** using *LIMON_IndEdges*. These interactions and corresponding edge importance obtained from LIONNESS are stored in the sample data frame along with the network statistics.

#### A5. Individual Network Edge Statistical Inference

The single sample network data is stored as a matrix per sample that includes the time points, subjects, covariates, centralities, and microbial taxa interactions for every sample. The *LIMON_NodeStat* function can then be used to conduct a linear regression for a continuous outcome, logistic regression for binary, or multinomial model for categorical outcomes, to identify which microbial interactions are associated with the users feature of interest (e.g., disease group, treatment arm, or continuous variables like blood sugar levels). Each model is run per unique interaction per time point. We set a minimum number of observations required per model type; 10 for linear regression, 20 for logistic regression, and 30 for multinomial. If an interaction at a time point does not have enough observations for model fitting, it is skipped and reported to the user. An adjusted p-value is calculated using the Benjamini-Hochberg correction (38). The model estimates, p-values, adjusted p-values, model type, number of observations per model, are all returned per interaction per timepoint. Users have the option to return a plot of edges that are statistically associated with their feature of interest by time for linear or logistic regressions. Plotting findings from the multinomial models is outlined in the package tutorial.

The full R package and tutorial can be downloaded from GitHub at https://github.com/LabBea/LIMON.

### B. Model Validation

#### B1. Synthetic Data Generation

Assuming that temporal dynamics of microbial communities can be modeled using the generalized Lotka-Volterra (GLV) equation, we generate synthetic data using the *miaSim* package in R (39):

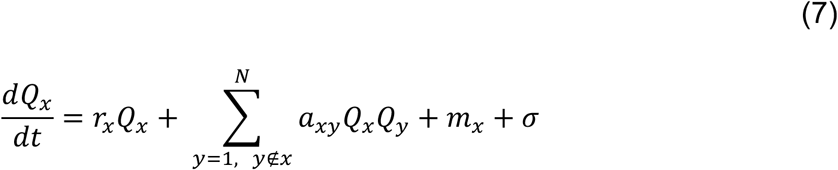

where *Q*_*x*_ represents the abundance of species *x*; *r*_*x*_ is the growth rate for species *x*; *y*_*xy*_ is the interaction value between species *x* and *y*; *m*_*x*_ is the migration rate (from/into the system); and *σ* is the model noise. For the general procedure, counts were simulated with a random initial covariance matrix by randomly sampling from a uniform distribution that was common across all the subjects. The default probability and deviation for migration rates were used (*m*=0.01). Simulated data were rounded, rescaled to be above 0 and multiplied by 10 to better resemble counts in real samples. Transformed simulated data follows a normal distribution (**Fig. S1A**) and produced variation in growth among the species (**Fig. S1B**). A covariate effect was added to the transformed simulated data by fitting a linear model:

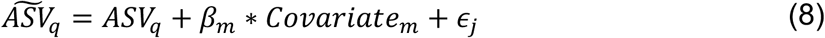

where 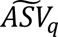 is the count value for species *q* accounting for the effect of covariate *m* with an effect of size *β*_*m*_, and ∈_j_, is an error term that follows a random uniform distribution.

To test the effects of different features in the performance of LIMÓN, we created 5 different datasets. First, we generated an initial dataset (Dataset 1, **Fig. 2**) using the approach described above and the resulting counts were inflated by an artificial binary covariate (to represent a binary variable such as sex, geographical location, or diet type) by randomly assigning the subjects to a covariate value of 0 or 1 and applying eq. 8 to ten of the fifty simulated taxa. After fitting, the values were rounded back and again inflated by a factor of ten to bring them to a similar order of magnitude than microbial sequencing results. From this general simulation, 4 additional sets of synthetic counts were simulated (**Fig. 2**, **S1 Table**): Dataset 2, sensitivity to number of subjects (subjects= 10, 20, 50, 75 and 100) ; Dataset 3, sensitivity to number of taxa (taxa= 10, 20, 50, 75 and 100); Dataset 4, sensitivity to network sparsity (sparsity level [% connectance]=0.1, 0.2, 0.5, 0.75 and 0.9); and Dataset 5, sensitivity to covariate strength (covariate strength= 0.01, 0.1,1,8, 15).

**Figure 2.**
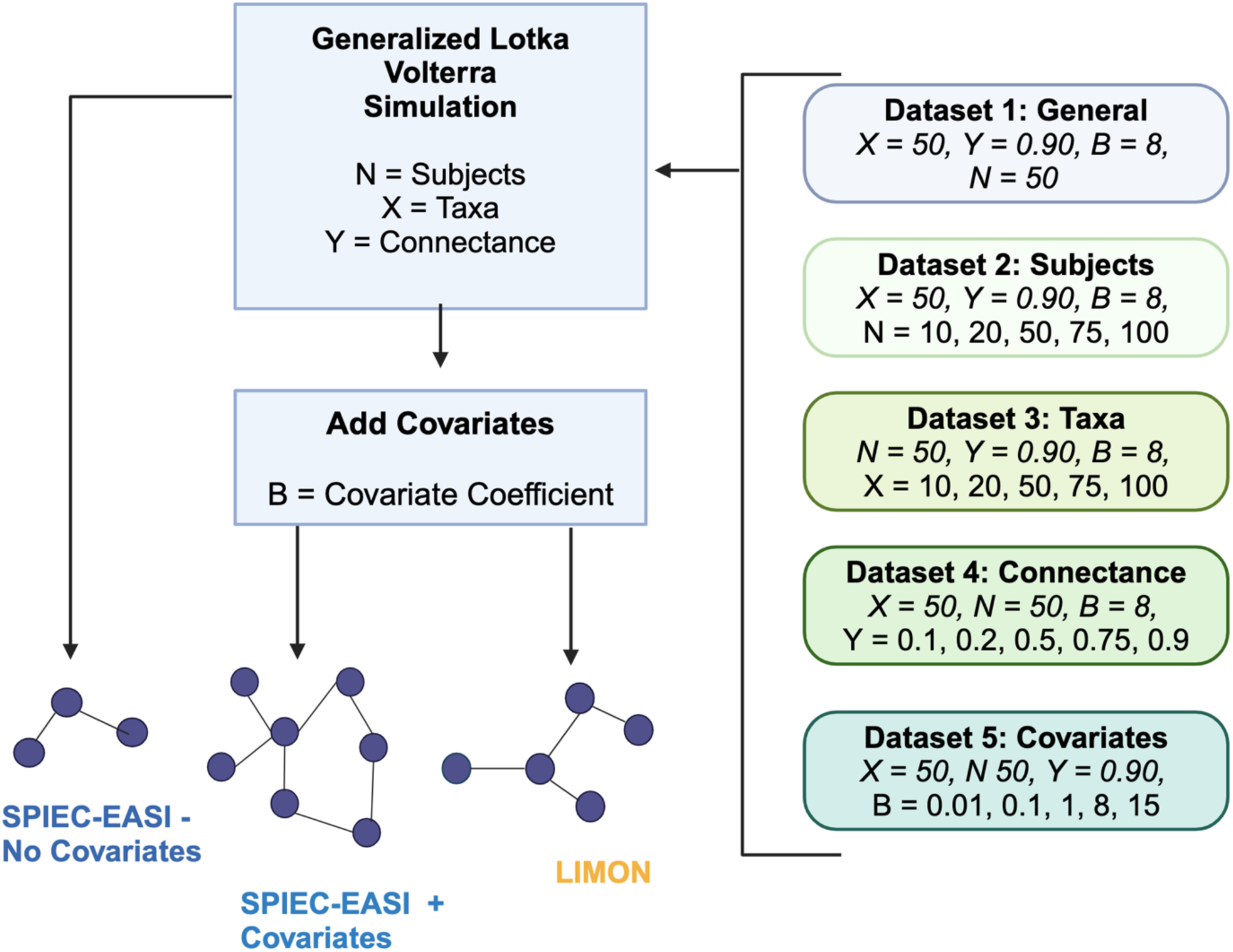
Sensitivity Analysis Workflow. A total of 21 datasets were generated using the generalized Lotka-Volterra (GLV) model and covariates were added to the simulated GLV data. For each dataset, co-abundance microbial networks were constructed using LIMÓN and SPIEC-EASI and contrasted against their correspondent reference networks (SPIEC-EASI no-covariates) that were created using GLV simulated data. Created in https://BioRender.com.

#### B3. Testing Single Sample Estimation Accuracy

To test whether the LIONESS procedure could return single sample estimates with microbial networks derived from SPIEC-EASI (the basis of LIMÓN), we created synthetic communities with differing numbers of subjects and species using the SPIEC-EASI data simulation function. We used GLV-generated counts generated above (section **B2**), and a common underlying graph structure in the SPIEC-EASI *synth_comm_from_counts()* function (**S2 Table, Fig. S2**). A zero-inflated negative binomial distribution was specified for these simulations. For each synthetic community, an overall network was estimated with SPIEC-EASI and we set to 0 the abundance of one of the species that was part of the strongest edge of the inferred network in one subject. Then we employed the LIONESS equation using SPIEC-EASI networks, instead of the package function with the default Pearson networks, to both the unaltered and altered count tables to return their individualized networks. We then compared the differences in edge per interaction per subject and extracted the top 10 edges with largest absolute difference.

Next, we explored whether LIMÓN was capable to simultaneously identified changes in edge importance in multiple interactions as a function of the number of subjects. For that, we identified the top five strongest non-overlapping interactions (all edges involved different species) from the adjacency matrix used in the SPIEC-EASI *synth_comm_from_counts()* simulation. We generated the basic count table from the simulation as above. From this, we created a total of 4 count tables by setting the value of one species from 3 or 5 of those tops edges to 0 in just 1 or 5 total subjects. Both the altered and unaltered datasets were taken through the LIONESS procedure with SPIEC-EASI network inference as above. The extracted edge tables were used to calculate the difference in edge value between the altered and unaltered data across the interactions that were changed (**Fig. S2B**).

#### B4. Covariate Removal Accuracy

To test if covariates can be removed from microbial counts while maintaining the longitudinal underlying nature of the community abundances, we removed the covariate effects from the GLV simulated data using *LIMON_DistrFit()* (**S3 Fig.**). We generated a simulated temporal count matrix of 50 taxa, 50 subjects, and 10 time points with a 0.9 connectance (Dataset 1, **Fig. 2**). We randomly selected two species, one with no covariate effect added (species A) and one with a covariate effect added by a linear model (species B). We used a linear mixed effect model in the *LIMON_DistrFit()* function and fit a covariate removal model for one species at a time. The denoised counts for both species A (should remain unchanged as no covariate effect was added) and B (should change with removal of the binary covariate) were extracted and compared to their respective original value from the simulation. Comparisons among the cases (no covariates data, covariates added, covariates removed) was performed using the Frobenius norm (40) of the covariance matrices obtained with Spearman correlations (**S2 Fig.**)

#### B5. LIMÓN Sensitivity to Detect Strong Interactions

Subsequently, we performed a sensitivity analysis to test LIMÓN’s robustness upon changes in the number of subjects, and taxa, the degree of network sparsity, and the covariance variability (Datasets 2- 5, see **Fig. 2** procedure). For each GLV generated dataset, we first inferred their co-abundance networks at each time point using SPIEC- EASI (*lambda* = 200, *lambda.min.ratio* = 0.01, absolute edge value threshold=0.0002). The large lambda, the graphical lasso regularization parameter, was chosen to increase sparsity and return strong edges. GLV simulated data inflated with the corresponding covariates values were input into SPIEC-EASI alone and LIMÓN to render a network per time point, per method (**Fig. 2**). The number of time points were maintained fixed to ten per sub-dataset/network inference method. We used the SPIEC-EASI co-abundance network returned from the GLV simulated data without the addition of covariates as our “*ground truth*”. Then we compared this ground truth network to those co-abundance networks obtained from the SPIEC-EASI and LIMÓN using covariate-adjusted data with the following: (1) the percent of *“ground truth”* edges recovered; (2) the number of total edges recovered by SPIEC-EASI and LIMÓN compared to the ground truth total; (3) the percentage of each edge weight category strength (low, medium and high from spearman correlation of the original GLV data). To establish the statistically significant differences between the percentage of *“ground truth”* edges recovered and the total number of edges as a function the different parameters (i.e. sample size, taxa size etc.) we employed the Wilcoxon rank sum test and corrected for multiple comparison with by Benjamin-Hochberg method (38).

#### B6. LIMÓN Recovery of individual network characteristics

One of the major advantages of LIMÓN is its capability to generate individual-specific and sample-specific co-abundance networks and return network characteristics per sample. To test this LIMÓN functionality, we generated sample-specific co-abundance networks in LIMÓN (50 subjects, 10 time points = total of 500 count matrices based on Dataset 1) and calculated, their individual network characteristics, such as node centrality, betweenness, and closeness. We then compared these network properties with (a) those of sample-specific networks created by LIMÓN if we did not remove the covariate effect; (b) and if we used the ground truth GLV simulated data (no covariate effect added). To compare to the ground truth, all centrality measures were averaged across the 10 simulated time points. Wilcoxon-rank sum with Benjamini-Hochberg p-value adjustment for multiple comparisons was utilized to compare differences in mean centrality measures among the three procedures.

### C. Model Application – Infant Microbiome Dataset

To show the utility of LIMÓN to detect individual microbial network changes over time in key outcomes of interest, we applied our pipeline to a publicly available longitudinal infant microbiome dataset involving a randomized control trial of infant diet during the first year of life (41). For this study, we used a total of 52 infants that were randomized into three dietary groups and had their fecal samples collected at 2, 4, 6, and 12 months of age (208 total samples). Diet types were normal breastmilk (BF), standard milk formula (SF) and an experimental formula supplemented with bovine milk fat globule membrane added (EF). Gut microbial species were profiled using 16S rRNA amplicon sequencing. Data for this study was extracted from the Curated Gut Microbiome and Metabolome Data Resource (42). For use in LIMON, we filtered to the top 75 most abundant taxa at the genus level. We normalized the raw counts using cumulative sum scaling. We then used LIMÓN to adjust for infant gender and render individualized networks per infant at all four time points, specifying a zero-inflated negative binomial model due to the high percentage of 0’s and overdispersion of the data. We then used LIMÓN to identify significant diet related microbial interactions at each monthly mile stone using the *LIMON_NodeStat()* function multinomial model with dietary group (BF, SF, EF) as the outcome. We chose the most significant interaction altered among the groups and characterized its difference by diet at each timepoint. To determine if this interaction could be linked to certain diets, the probability and confidence intervals for a range of edge weights for this interaction were estimated using R *ggeffects* package. The importance of that edge to the model prediction of diet was compared to a null model (intercept only) using a likelihood ratio test from the R *car* package.

### D. Data Availability

All the scripts to simulate and test each set of data can be found on the project GitHub at https://github.com/LabBea/LIMON_Validation.

## RESULTS

### Single Sample Estimates can be derived for Microbial Networks

First, we tested whether LIONESS was able to recover individualize co-abundance microbial networks when using SPIEC-EASI to determine the network structure (**Fig. 3**). When the count data of a strongly connected species was set to 0 in just one subject at random (species not present in the community), the co-abundance network of that given subject with that species presented the largest number of differences when comparing the generated co-abundance networks with the unaltered and altered data. The strongest results were seen when 100 taxa were included in the network estimation rather than 50 (**Fig. 3A**). We further tested whether LIONESS with SPIEC-EASI could recover individualized co-abundance networks when multiple subjects lost more than one highly connected species. We set the edges of one species in 3 or 5 strongest edges to 0 in 1 or 5 subjects and found the largest changes in edge values were in those subjects/species (**Fig. 3B**). Our results indicates that LIONESS when using SPIEC-EASI as underlying model can adequately detect individualized changes in co-abundance microbial networks but performance improved with a higher ratio of taxa to samples (50 subjects, 100 taxa; 100 subjects, 100 taxa).

**Fig 3:**
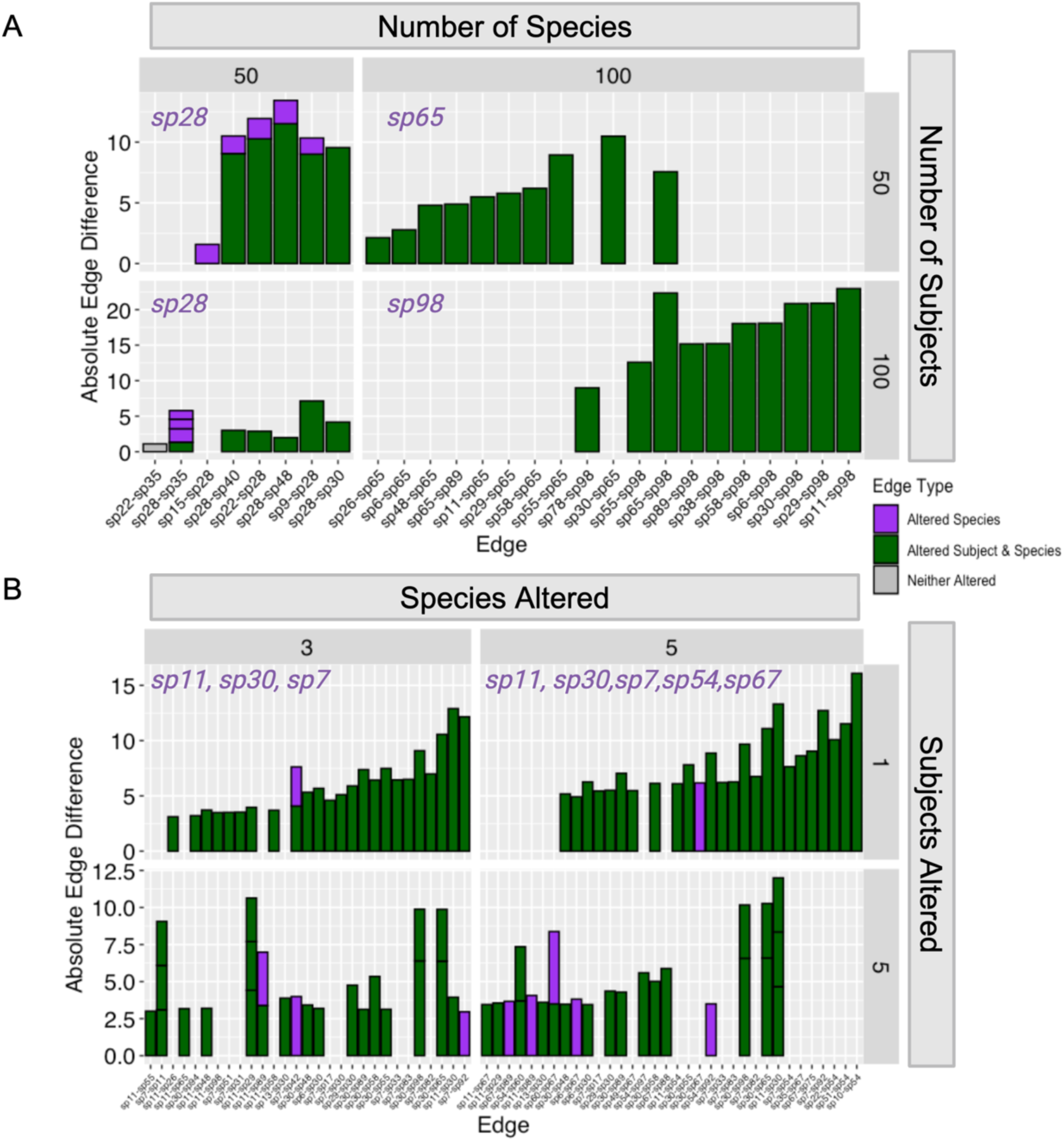
LIONESS/SPIEC-EASI Identifies Single-Sample Alterations in Microbial Networks. (A) The top 10 most different edges after removing one highly connected species from a microbial community. (B) The top 25 most different edges after removing three or five highly connected species from the community in one or five individuals. Green indicates that the edge was present in the individualized co-abundance network of the altered subject and was part of the interaction pattern of the removed species. Purple indicates that the edge was part of the interaction pattern of the removed species but was found in the individualized co-abundance network of a non-altered subject. Grey indicates that neither the species nor the subject had any data altered. Each box in the bar represents an absolute edge difference value. Species altered in each iteration are annotated in purple.

### LIMÓN Removes Covariate effects from Microbial Data

Then we explored whether LIMÓN could remove the effects of subject covariates (e.g., age, weight, sex) from in temporal microbial communities (**Fig. S4**). For that, we inputted into LIMÓN a simulated count dataset of 50 taxa and 50 individuals generated with the GLV model and the same set but inflating the GLV counts with a binary term to simulate covariate variability (see Methods for more detail). Species that did not have a covariate effect (species A) applied to them showed only a small deviation in their counts after undergoing the covariate removal step in LIMÓN (**S4B Fig.,** sd=2.5). Simulated species that were inflated with a binary covariate effect (species B) exhibited a shift in its counts when compared to the simulated GLV data with no covariate effect (**S4C Fig.,** sd = 40.3). LIMÓN, as expected, removed the effect of the binary covariate (**S4D Fig.**, sd=5.8). In fact, the covariance matrices of the GLV simulated data and of the LIMÓN adjusted counts were very similar (**S6 Fig.**, Frobenius norm=0.53) In summary, LIMÓN successful removes covariate effects in sparse count data and while maintaining the underlaying temporal associations between the members of the microbial communities.

### LIMÓN retrieved more accurate edges than SPIEC-EASI using covariate conflated GLV-simulated data

Recovery of the “*ground truth*” network structure from covariate conflated data revealed that LIMÓN outperformed SPIEC-EASI across multiple scenarios. When provided temporal count data generated using the GLV model that were covariate conflated to LIMÓN (provides covariate correction) or to SPIEC-EASI (no covariate correction), LIMÓN recovered a higher percentage of ”*ground truth*” edges with lower number of false positive edges with increasing sample size, (**Fig. 4A, B**), microbial community diversity (number of taxa, **Fig. 4C, D**), microbial community complexity (connectance, **Fig. 4E, F**) and, unsurprisingly, with noise (**Fig. 4G, H**). SPIEC-EASI on covariate conflated-data consistently returned more spurious connections with increasing sample size, strength of covariates, and number of taxa. Taken together, these results demonstrate LIMÓN can better return underlying microbial interactions even in small size longitudinal experiments while reducing spurious interactions caused by confounding factors.

**Fig 4.**
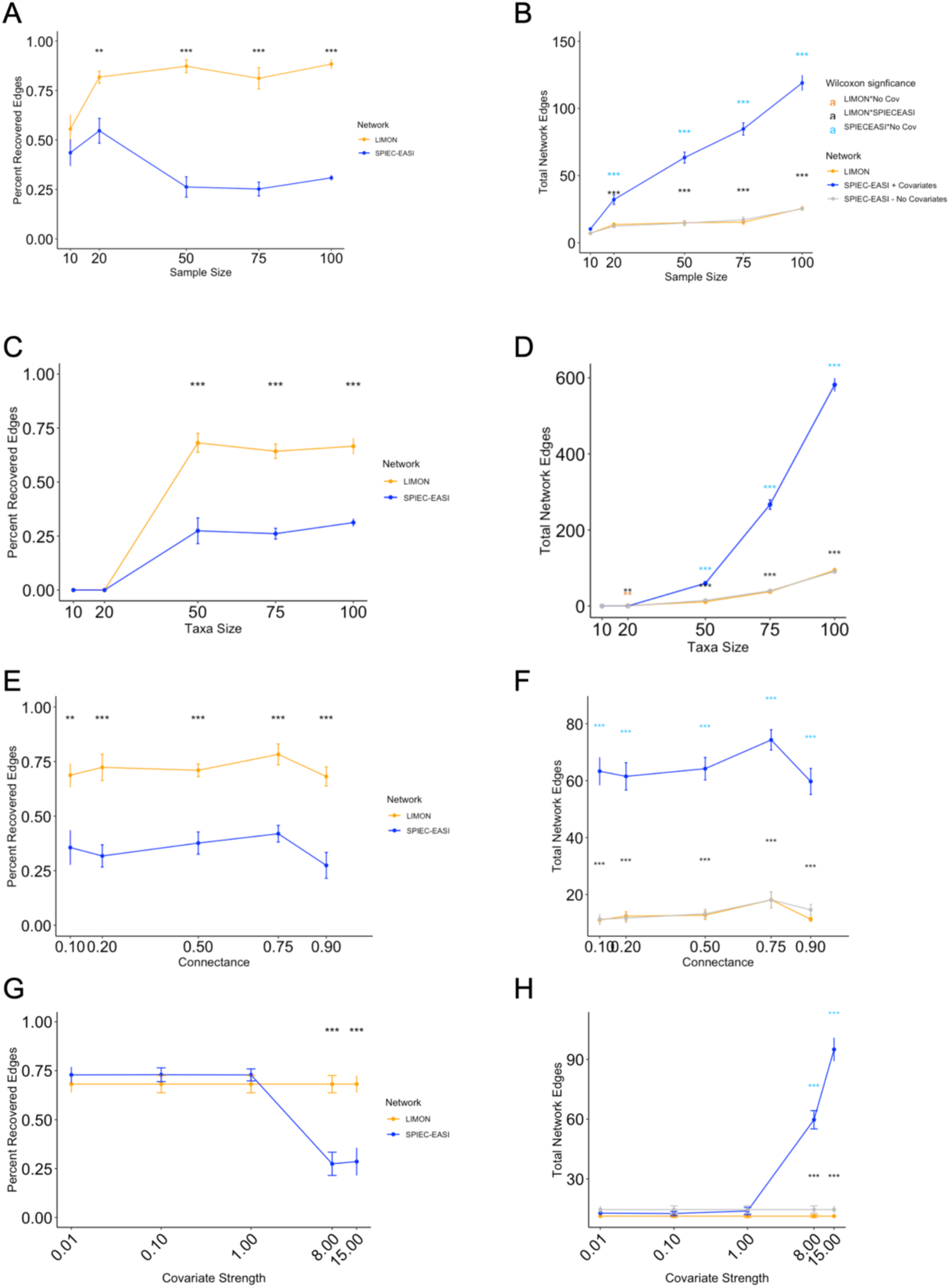
LIMÓN recovers more true edges and fewer false positive edges than SPIEC-EASI alone. Sensitivity to the sample size (number of subjects, dataset 2) in terms of percentage recovery edges (A) and total number of network edges (B) compared to those recovered by SPIEC-EASI using the raw simulated count data generated using the GLV model. (C & D) Sensitivity to microbial diversity (number of taxa, dataset 3); (E & F) Sensitivity to community complexity (sparsity of connections,, dataset 4); (G & H) Sensitivity to noise level in the data (covariate coefficient strength, dataset 5) Wilcoxon rank sum test adjusted p-values for multiple comparisons (Benjamini-Hochberg): NA = not significant, *p < 0.05, **p < 0.01, ***p < 0.001. Plot G & H values were plotted on a log x-axis for clarity.

### LIMÓN recovered ground truth individualized network characteristics

LIMÓN better returned the “*ground truth*” network characteristics (co-abundance network inferred with SPIEC-EASI on no covariate data”) such as node degree, number of communities, closeness, betweenness, and eigenvector centrality compared to SPIEC-EASI with covariate effects added (**Fig. 5**). While the mean centralities among the three groups were all significantly different (p < 0.001), network properties generated by LIMÓN on covariate-conflated data closely resembled the ground truth. In contrast, SPIEC-EASI co-abundance networks, created on covariate-conflated data without adjusting for subject characteristics, exhibited more spurious and false-positive connections, as indicated by higher node degree and eigenvector centrality. These results further support LIMÓN’s effectiveness at returning underlying networks structure despite the presence of confounding factors in the raw count data.

**Fig 5.**
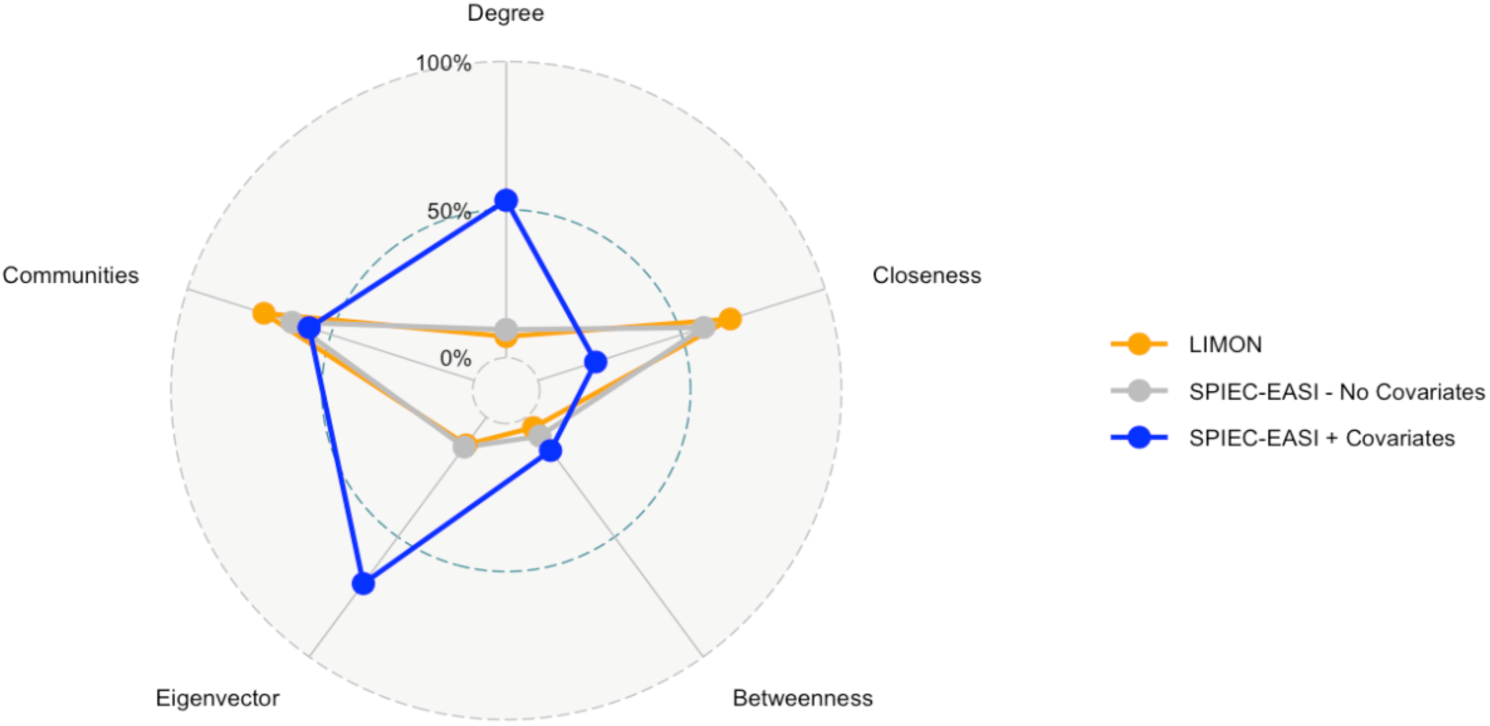
LIMÓN accurately recovered the network characteristics of the GLV-simulated data than SPIEC-EASI in the presence of conflated data due to differences in subject characteristics. Data were scaled between 0 and 1 by centrality measure and then averaged by method. Centrality measures of betweenness, closeness, number of communities, eigenvector, and node degree were normalized between 0 and 1 and compared using pentagon plots. All centralities compared between methods with Wilcoxon Rank Sum were significant (p < 0.001).

### LIMÓN identifies novel temporal changes in infant gut microbial communities

To demonstrate the utility of LIMÓN when studying longitudinal cohort data, we used a publicly available dataset of 52 infants that were randomly assigned to three different diets; standard formula (SF), bovine protein supplemented experimental formula (EF) or breast milk (BF), and whose gut microbiome was profiled at 2, 4, 6 and 12 months of age (**Fig. 6A**). He and colleagues hypothesized there would be differences in the gut microbiome and fecal metabolome between infants who received SF vs BF, and that this difference would be mitigated in infants who received EF instead of SF (41). Using LIMÓN, we identified 24 edges that were significantly different among the three dietary groups at different stages of development (2, 4, 6 and 12 months). The most significant edge was an interaction between the genus *Parabacteroides* and *Actinomyces* at 6 months. This interaction was not present in enough samples at 2 and 4 months to fit a multinomial model, thus we compared the 6 month and 12-month interactions. At 6 months, this interaction was able to distinguish among the three dietary groups when g_*Parabacteroides* and g_*Actinomyces* had positive edge weight (positive interaction) indicating a higher probability of the infant consuming an SF diet (**Fig. 6B**, 66% probability of SF, 30% of EF, and 4% of BF at an edge weight of 2.5). However, this distinction was no longer present at 12 months (**Fig. 6C**). A likelihood ratio test comparing an intercept-only model showed that the *g_Parabacteroides*–*g_Actinomyces* edge weight was a significant predictor of diet type at 6 months (p < 0.05), but not at 12 months (p > 0.05). LIMÓN was able to go beyond traditional single species identification and render differentially abundant community interaction associated with the differing diet types across the first year of life, while also controlling for infant sex.

**Figure 6.**
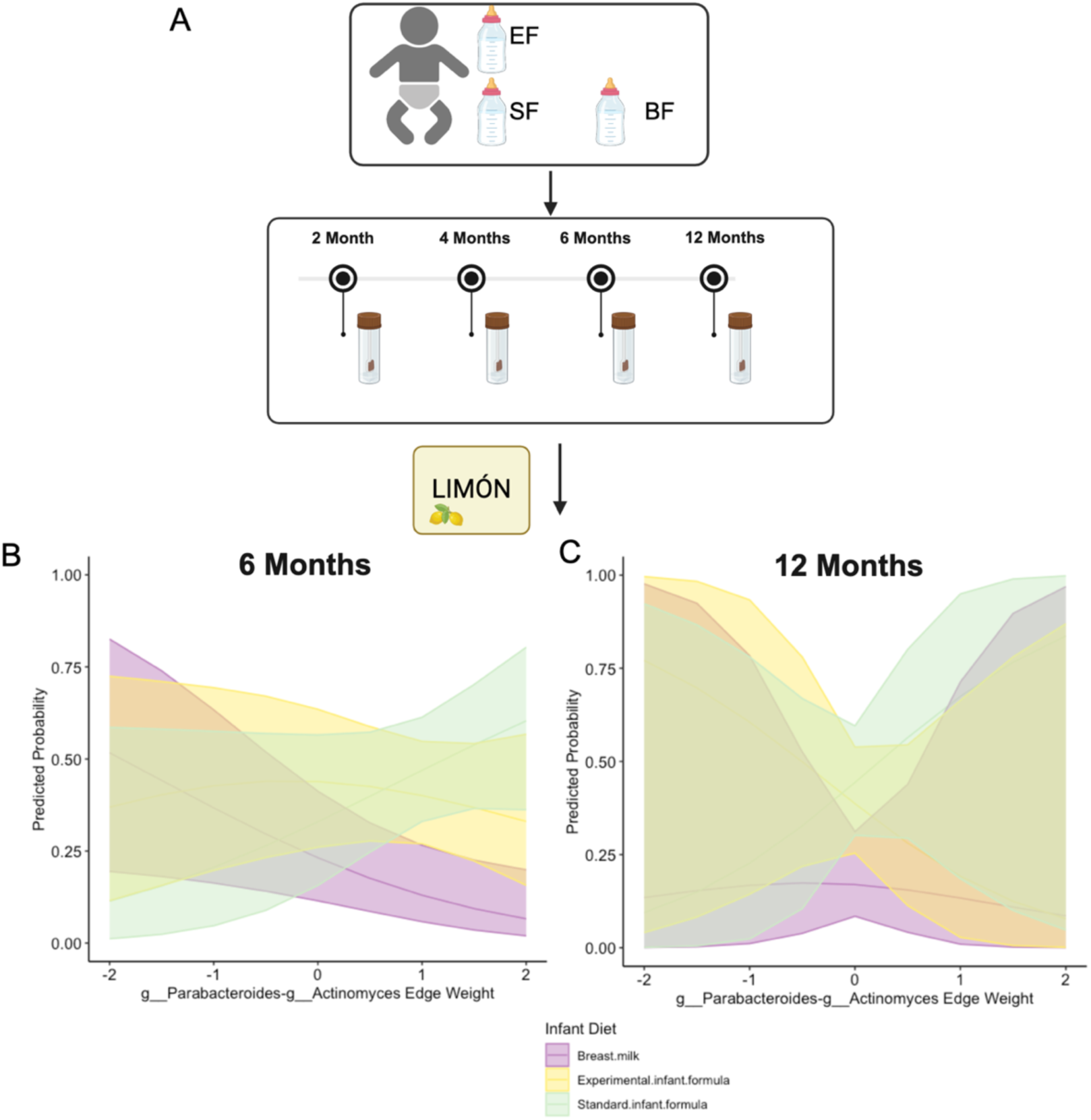
LIMON identifies diet specific differentially abundant network interactions overtime. (A) Experimental design. Infants were randomly assigned to one of three dietary groups; SF = standard formula, EM = Bovine protein supplemented experimental formula, BF = breast fed. Gut microbiome and fecal metabolome was sampled at 2, 4, 6, and 12 months. Microbial count data was taken through the LIMON procedure. (B) Probability of infant diet based on edge weight value between the genus *Parabacteroides* and *Actinomyces* at 6 months. A likelihood ratio test comparing an intercept-only model showed that the *g_Parabacteroides*–*g_Actinomyces* edge weight was a significant predictor of diet type at 6 months (p < 0.05). (C) Probability of infant diet based on edge weight value between the genus *Parabacteroides* and *Actinomyces* at 12 months. A likelihood ratio test comparing an intercept-only model showed that the *g_Parabacteroides*–*g_Actinomyces* edge weight was not a significant predictor of diet type at 6 months (p > 0.05).

## DISCUSSION

LIMÓN is an open-source R pipeline that enable researchers to infer individualized co-abundance microbial networks from dynamic processes. As microorganism do not work in isolation, it is essential to take a systems approach to investigating these communities that can reveal the complex interactions between the elements in the microbial communities. By providing multiple networks properties at the individual sample level (edge weight, centrality measures), LIMÓN also enables researchers to identify the keystone microorganisms and their connections by computing community features to a key outcome of interest.

Our results demonstrated that LIMÓN overcomes most limitations of current methods for capturing complex interactions of microbial communities in dynamic processes (e.g., disease progression, infant development), such as the need for removal of the effect of covariates in the spread of the data (most common in human studies), adequate methods to handle sparse compositional data, small sample sizes, and, most relevant for precision medicine, building individualized networks. With precision medicine/science becoming a more prominent focus in research, having a tool like LIMÓN that can provide individualized network estimates becomes even more integral to microbial research.

By leveraging mixed effects models and two existing tools, SPIEC-EASI and LIONESS, LIMÓN can construct individual longitudinal networks. First, we have shown that LIMÓN could remove covariate effects from repeated measures using mixed effect models designed for microbial compositionality, even with small sample sizes (**Fig. 4**). The small shift in values after covariate removal for taxa without covariate effects is likely due to the estimated effect for those taxa still being removed. Since LIMÓN was still able to accurately recover the underlying network after covariate removal, these shifts were considered negligible in terms of network structure. However, future versions of LIMÓN will include testing a p-value threshold to remove covariate effects only for taxa where they are significant. LIMÓN returned the underlying network structure with a lower false discovery rate compared to SPIEC-EASI alone when tested on data containing confounding effects, a given in real world studies. These sensitivity analysis findings indicate LIMÓN is more robust to changes in community complexity than SPIEC-EASI alone. Failing to removal of variables that are known to affect the microbial community (e.g., sex, weight, age) leads to complex networks fill with spurious connections (**Fig. 4**), thus preventing the identification of the true underlaying microbial communities. Likewise, estimation of the individualized networks illustrates how network characteristics per sample (e.g., centrality, number of communities) were better characterized by LIMÓN’s procedure than producing single sample networks from SPIEC-EASI alone. This feature of LIMÓN, the correction for covariates, is essential when human cohort studies. MAGMA is a new approach that also aims to remove the effect of covariates in microbial count data using an elegant combination of zero-inflated negative binomial models and Copula Gaussian graphical models. However, MAGMA cannot adjust time-series microbial data. Second, we have demonstrated that the simple LIONESS linear interpolation between the covariance matrices of SPIEC-EASI co-abundance microbial networks implemented in LIMÓN can discriminate individual microbial community networks (**Fig. 3**). Construction of microbial co-abundance networks from cross-sectional and longitudinal cohort studies have been based on the average interactions per group or condition (case and control studies) or a given time point. Recent work integrated MAGMA and LIONESS to render individualized microbial networks in patients with irritable bowel disease. Howver LIONESS by defualt employs standard correlation inference techniques, such as Person or Spearman correlations to establish the associations between microbial species. Standard correlations have been shown to perform poorly with compositional and sparse data as they tend to produce spurious interactions (16,18). LIMÓN employs the covariance matrix produced by SPIEC-EASI, which was specifically developed for sparce microbial networks.

LIMÓN is a validated open-source package to generate and analyze individual dynamic community networks, even with small sample sizes (**Fig. 3A**). The key applicability of LIMÓN lies in its ability to perform differential analysis of community interactions or features rather than on independent species. In the published analysis of an infant dietary interventional study, He and colleges hypothesized that differences in the concentration of oligosaccharides vs proteins in infant formula would alter the gut microbiome across the first year of life (41). Using LIMÓN, we could go beyond single species identification, and determine which microbial interactions differed by dietary type over the first year of life. Specifically, we identified a key interaction between two common gut microbial genus, g_*Parabacteroides* and g_*Actinomyces,* was highly predictive of the standard formula diet at 6 months. Study authors had hypothesized that the experimental formula would mitigate some of the microbial and metabolic differences observed between infants who were breast fed vs standard formula fed. They found the formula groups were more similar to each other than BF at 2, 4 and 6 months, but indistinguishable from the BF group at 12 months. LIMÓN both supports and expands these findings. First, the association between these two microbes being strongly linked to SF may explain the metabolic observations of the study. *Parabacteroides* is a known producer of short chain fatty acids (43,44), which were higher in the formula groups at 6 months. Second, this interaction was strongest at the 6-month period, before becoming indistinguishable from the BF group at 12 months, same as what was observed at the alpha and beta diversity level in the original findings. Having a community interaction level understand of these gut communities may help better explain the dynamics related to the metabolic alterations better than single species or broad diversity measures alone. This is one example of how LIMÓN can provide a more complete understanding of how the microbial community dynamics change in cohort-based studies.

LIMÓN has several strengths, including estimating single sample networks, adjusting for confounding effects, and returning temporally meaningful community interactions. Further, LIMÓN is user-friendly with capabilities to extract intermediates and final datasets, retrieve network adjacent matrices and their centrality measurements per individual and time point, perform differential analysis of network features, and be easily export to other visualization platforms, such as Cytoscape (45). Additionally, the users have flexibility for choosing the appropriate mixed effect model distribution for their data, which can help ensure appropriate modeling for many data types (46). There are several areas for improvement that LIMÓN would benefit from. For instance, LIMÓN can account for continuous and binary covariate data but cannot yet handle multilevel categorical covariates nor missing data, if the sample is not present for that time point, no network will be estimated for it.

In future versions, we would like to incorporate methods to handle categorial variables as well as missing data from longitudinal microbial sequencing information (47). LIMÓN currently employed SPIEC-EASI to infer the microbial co-abundance network structure other methods are available to identify interactions in microbial communities. Incorporating more methods for network inference in the future can provide better flexibility users and to take advantage of ensemble methods. Further work into incorporating causal role of bacterial communities in disease states could also improve LIMÓN’s utility (48). Finally, LIMÓN might not detect subtle individual differences in microbial network structure as it employs linear interpolation. Other non-linear approaches might be better for those cases.

## CONCLUSION

Methods for capturing complex interactions of microbial communities are often limited due to the need for large sample sizes, their inability to handle longitudinal data or the effect of covariates in the spread of the data, and/or their inability to build individualized networks. LIMÓN can overcome most of these limitations by removing covariate effects and providing individualized estimations of microbial networks over time in a user-friendly R package. In doing so, LIMÓN opens a new avenue of microbial community precision science by producing individualized network descriptors to capture heterogeneity in diverse communities.

## DATA AVAILABILITY

LIMÓN is published as an R package. The package and a full tutorial can be found on GitHub at https://github.com/LabBea/LIMON. All scripts used to simulate and analyze data are reproducible and located at https://github.com/LabBea/LIMON_Validation.

## ACKNOWLEDGEMENTS

SAA and BPB conceptualized and designed the methodology for this work. SAA was responsible for the software, validation, and formal analysis. BPB provided supervision and funding acquisition. SAA and BPB wrote the manuscript. SAA was funded by the University of Illinois Medical Scientist Training Program. BPB was funded by BIRCWH K12 Award (K12AR084225)

