## Supplemental Figures for "Creation and validation of LIMÓN - Longitudinal Individual Microbial Omics Networks"

A

B

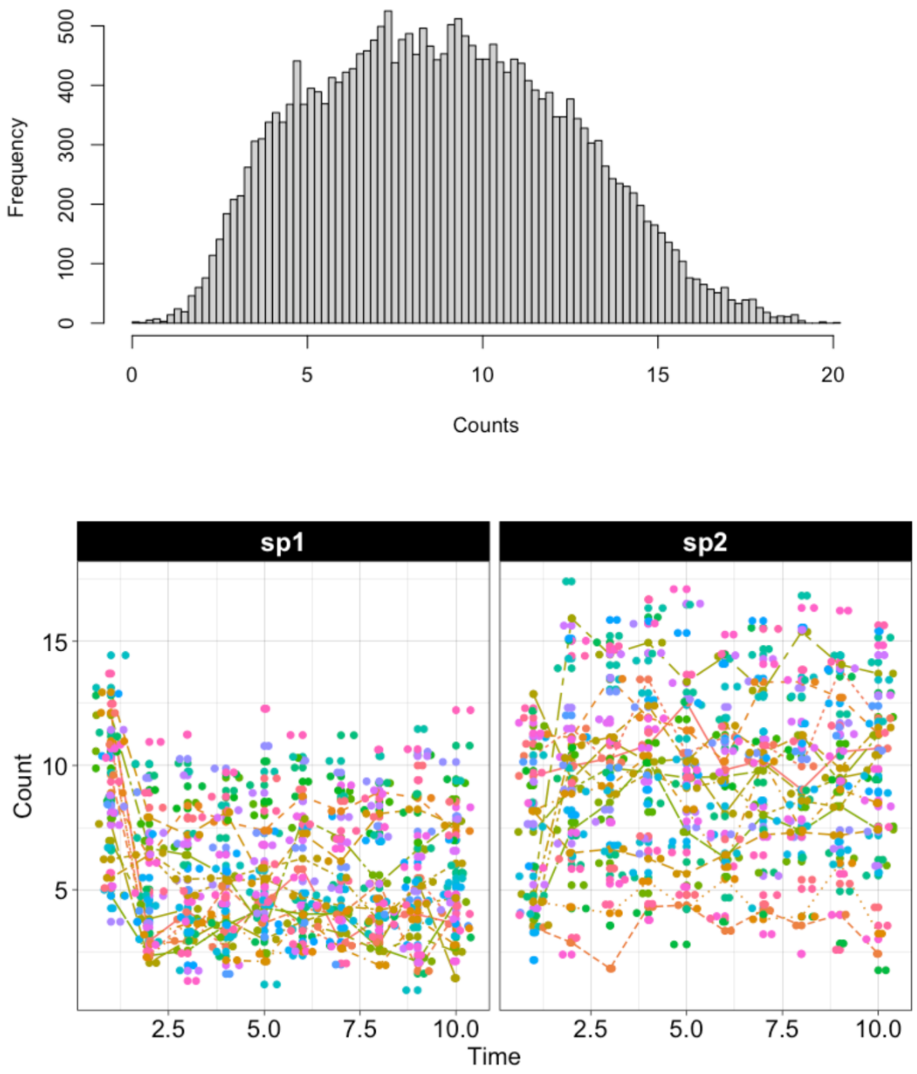


**Figure S1. Simulated data generated using a generalized Loka-Volterra (GLV) model.** (A) Distribution of the GLV simulated data using the parameters described in Dataset 1 (50 subjects, 50 Taxa, 90% connected). (B) Temporal trends of for two example taxa, species 1 (sp1) and species 2 (sp2) based on the GLV model. Colors represent different simulated subjects.

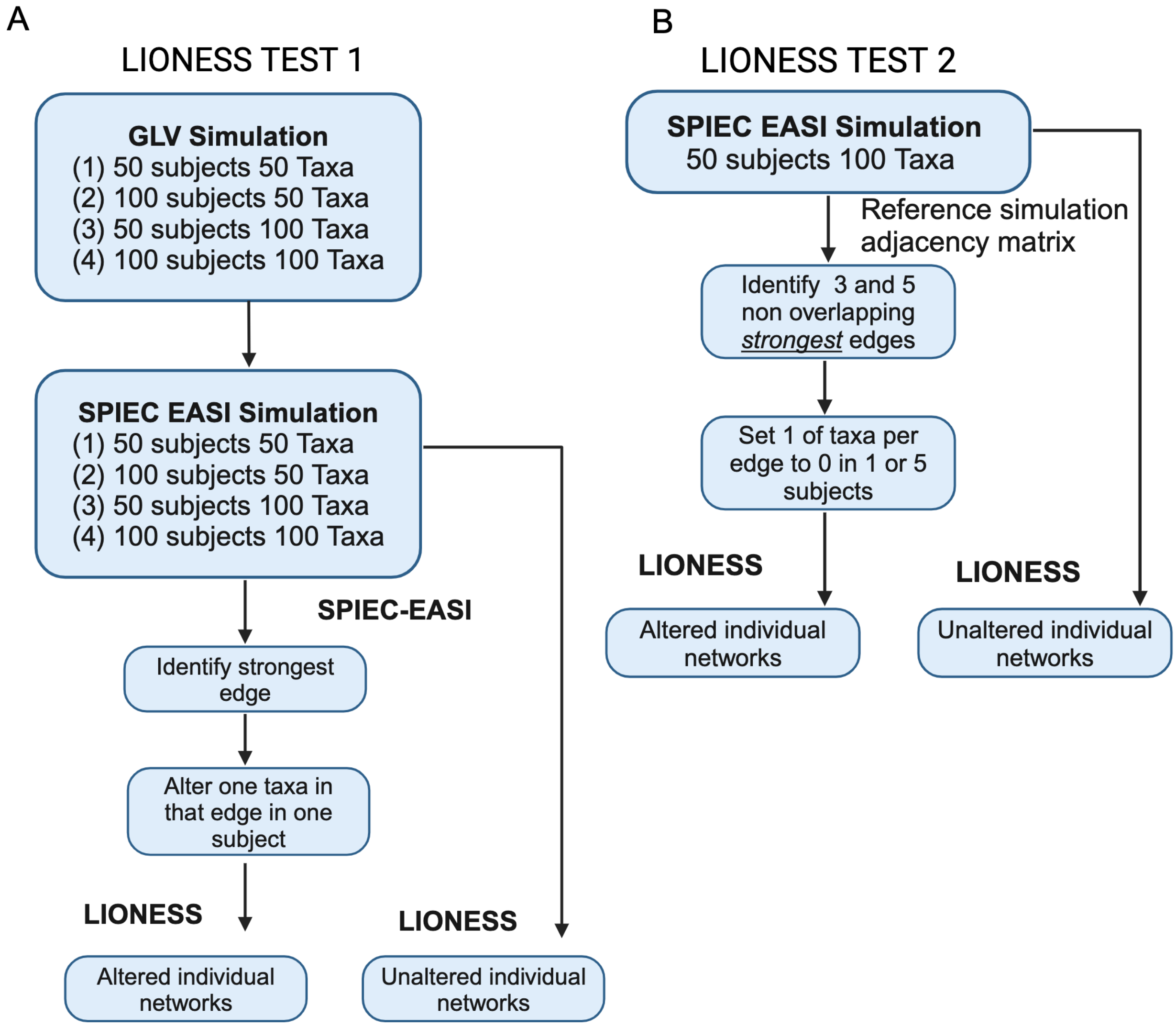


**Figure S2. Single Sample Estimation Testing Workflow. (A) LIONESS Test 1** – the value of one taxa in the SPIEC-EASI identified strongest edge was set to 0 in one subject. LIONESS procedure was completed for the altered and unaltered datasets and difference in edges were compared. **(B) LIONESS Test 2** – The 3 or 5 strongest non-overlapping edges were identified from the *synth_comm_from_counts()* simulation adjacency matrix. 1 taxa per edge was set to 0 in 1 or 5 subjects. Altered and unaltered data underwent the LIONESS procedure (with SPIEC-EASI), and edge differences compared. Figure created with BioRender.com.

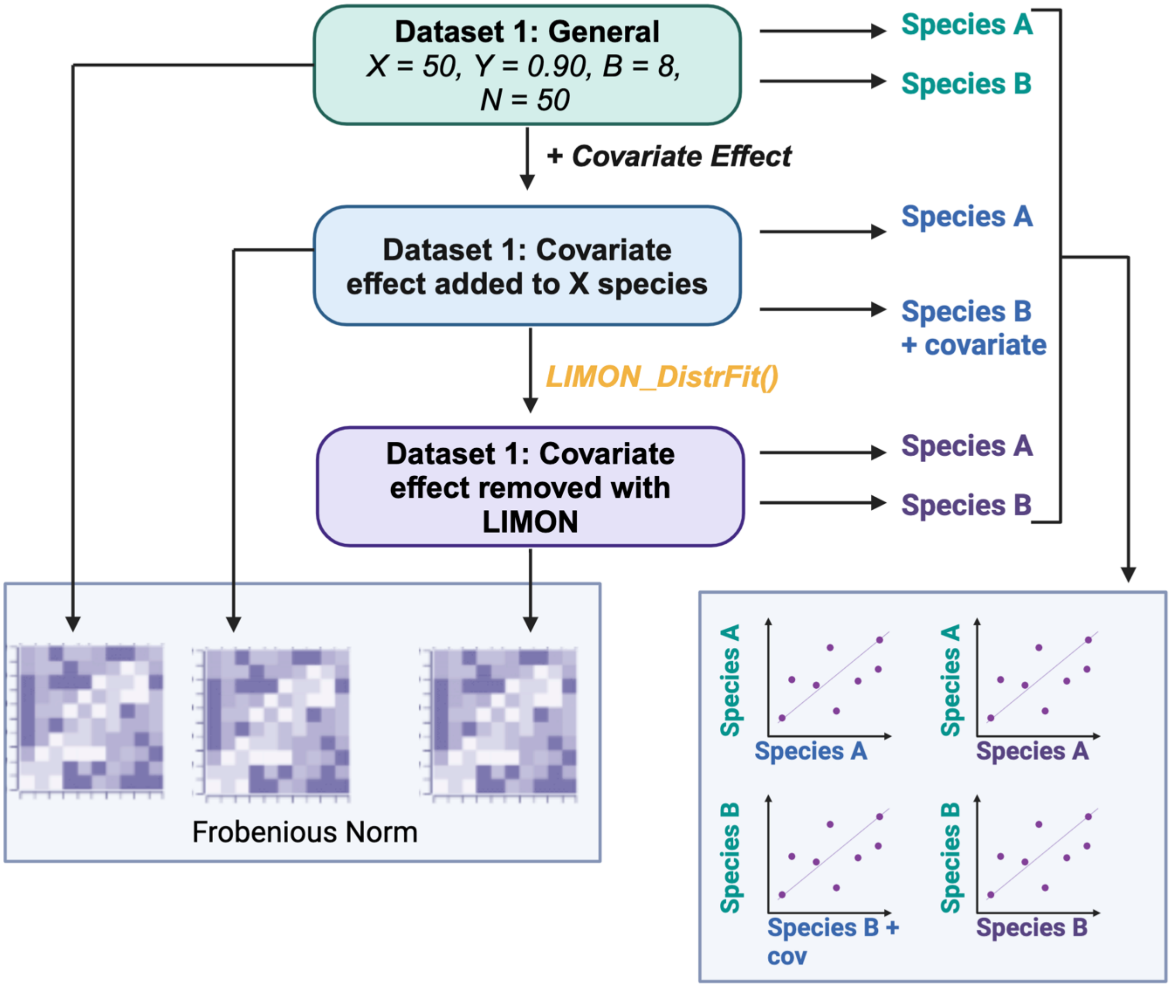


**Figure S3. Procedure for testing covariate removal in LIMON.** Data were simulated from the GLV using 50 subjects (X), 50 species (N), 90% connected species (Y), and a covariate effect size of 8. The covariate effect was added to 10 of the 50 species. The dataset was taken through *LIMON_DistrFit()* step specifying the binary covariate for removal. We compared the counts of the original simulated data (no covariates added, green), data with a covariate effect added (blue), and data after covariate effect removal with LIMÓN (purple). We compared the overall spearman correlations using the Frobenious norm and then compared the counts of two representative species. Species A had no covariate effect ever added, Species B had a covariate effect added. Figure created with BioRender.com.

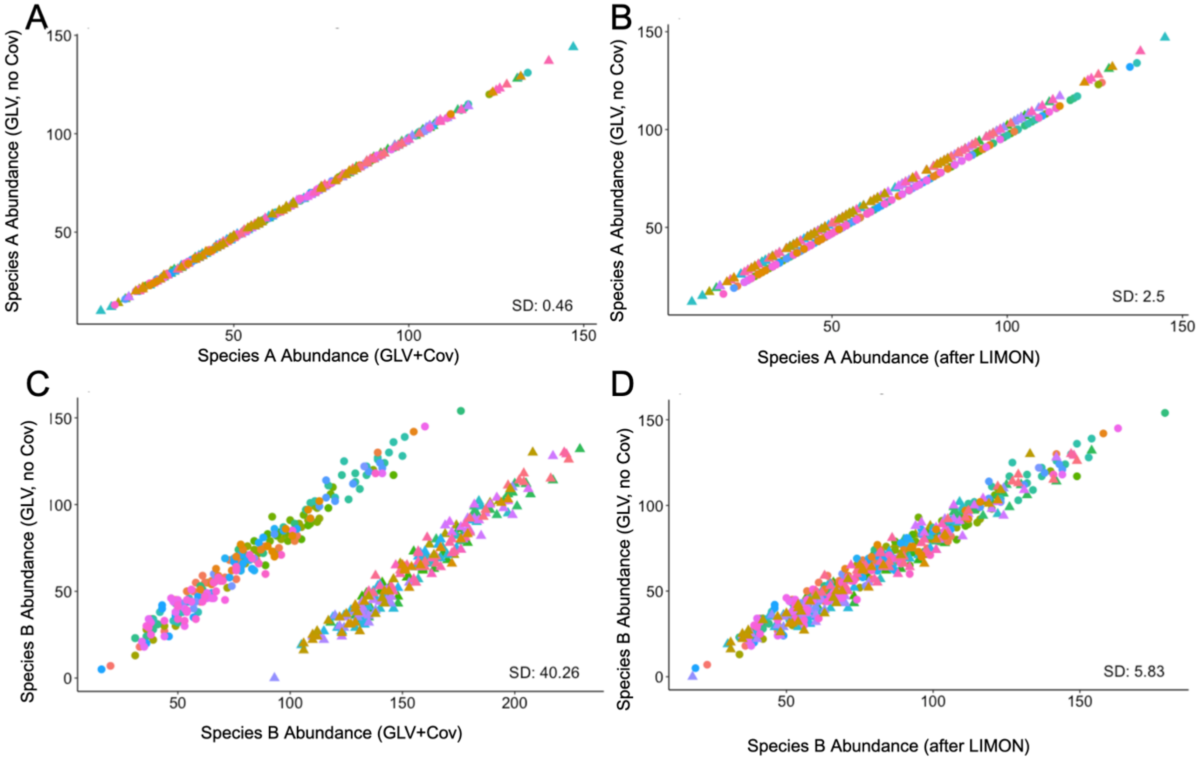


**S4. LIMÓN Removes Covariate Effects.** (A) Simulated GLV count with the covariate effect (x) versus the original simulated GLV count before covariate effect was added (y) for species A. Standard deviation = 0.46. (B) Species A transformed data after undergoing mixed effect correction in LIMÓN (x) compared to original simulated GLV count before covariate effect was added (y). Standard deviation = 2.5. (C) Simulated GLV count with the covariate effect (x) versus the original simulated GLV count before covariate effect was added (y) for species B. Standard deviation = 40.26. (D) Species B transformed data after undergoing mixed effect correction in LIMÓN (x) compared to original simulated GLV count before covariate effect was added (y). Standard deviation = 5.83. Points are colored by simulated subject ID; each ID has 10 timepoints. All data were generated using the following parameters (Dataset 1): 50 taxa (X) and 50 subjects (N), with a covariate coefficient of 8 and a connectance value of 0.9. Diamonds are covariate = 0, circles are covariate = 1.

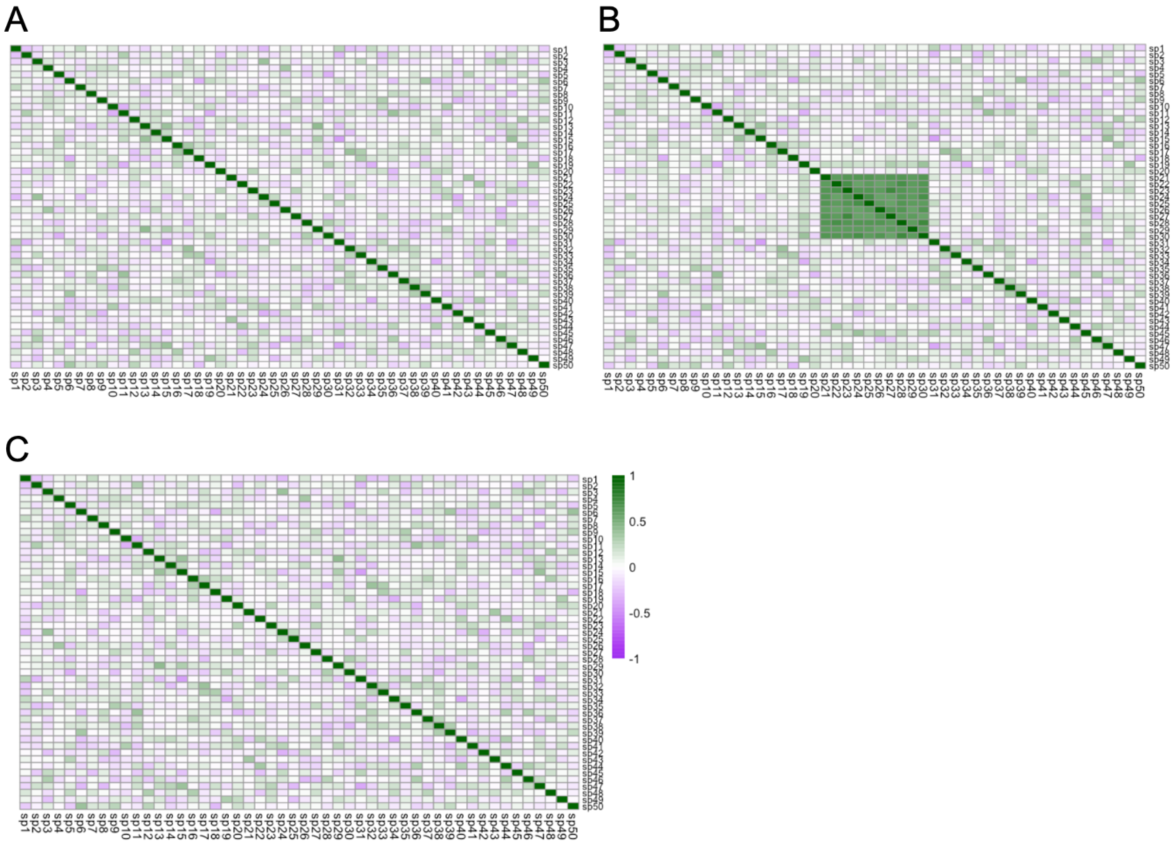


**Figure S5. LIMÓN corrected counts reflect simulated GLV counts.** (A) Spearman correlation of counts before addition of covariates to Dataset 1 (50 taxa (X) and 50 subjects (N), with a covariate coefficient of 8 (B) and a connectance value of 0.9 (Y)). (B) Spearman correlation of counts after addition of a binary covariate to simulated species 21-30, this is the increased connectance in green. (C) Spearman correlation of LIMÓN corrected data from Dataset 1. Frobenius norm of A – B is 6.8, and A – C is 0.53.
